# The Use of Fluid-phase 3D Printing to Pattern Alginate-gelatin Hydrogel Properties to Guide Cell Growth and Behaviour *In Vitro*

**DOI:** 10.1101/2023.07.08.547691

**Authors:** Souza Andrea, McCarthy Kevin, Rodriguez Brian J., Reynaud Emmanuel G

**Affiliations:** School of Biomolecular and Biomedical Science, University College Dublin, Belfield, Dublin 4, Ireland; School of Physics, University College Dublin, Belfield, Dublin 4, Ireland

**Author notes:** ^†^The author has died prior to the submission of this paper. This is one of his last works.

**Keywords:** 3D (bio)printing, embedding 3D (bio)printing, fluid-phase, patterning, alginate-gelatin, U2OS.

## Abstract

3D (bio)printing technology has boosted the advancement of the biomedical field. However, tissue engineering is in its infancy and (bio)printing biomimetic constructions for tissue formation *in vitro* is still a default. As a new methodology to improve *in vitro* studies, we suggest the use of a cross-linkable aqueous support bath to pattern the characteristics of the scaffolds during the 3D printing process. Using fluid-phase, different molecules can be added to specific locations of the substrate promoting cell behaviour guidance and compartmentalization. Moreover, mechanical aspects can be customized by changing the type or concentration of the solution in which the (bio)printing is acquired. In this study, we first assessed different formulations of alginate/gelatin to improve cell colonization in our printings. On formulations with lower gelatin content, the U2OS cells increased 2.83 times the cell growth. In addition, the alginate-gelatin hydrogel presented a good printability in both air and fluid-phase, however the fluid-phase printings showed better printing fidelity as it diminished the collapsing and the spreading of the hydrogel strand. Next, the fluid-phase methodology was used to guide cell colonization in our printings. First, different stiffness were created by crosslinking the hydrogel with different concentrations of CaCl_2_ during the printing process. As a result, the U2OS cells were compartmentalized on the stiffer parts of the printings. In addition, using fluid-phase to add RGD molecules to specific parts of the hydrogel has also promoted guidance on cell growth. Finally, our results showed that by combining stiffer alginate-gelatin hydrogel with RGD increasing concentrations we can create a synergetic effect and boost cell growth by up to 3.17-fold. This work presents a new printing process for tailoring multiple parameters in hydrogel substrates by using fluid-phase to generate a more faithful replication of the *in vivo* environment.

## 1. INTRODUCTION

Since the 1980s, 3D printing technology has become one of the most disruptive technologies in science, especially in the biomedical fields. For instance, 99% of the fitting hearing aids are produced by 3D printing. Moreover, spinal, hips, and dental implants are also important deliveries of the 3D printing technology [1]. However, more complex biomedical fields, such as tissue engineering and organ regeneration, are still in their infancy due to the complexity of the process involved in replicating tissue’s characteristics *in vitro*. In this case, multiple parameters are required to produce a faithful structure that would allow the success of this approach, such as 1) the choice of the biomaterial or bioink and possible combination of different materials; 2) the addition of different ECM and other molecules, such as growth factors, to the scaffold to modulate cell adhesion, proliferation, migration, and differentiation [2]; 3) the best 3D printing methodology to ensure cell viability and/or printability of the designed scaffolds; 4) the crosslinking process to secure the printing stability over time and to control the degradability rate; 5) the cell type and the seeding protocol. All these variables can modify the scaffold’s physical, chemical, and biological aspects, and defining the ideal protocol to reproduce *in vitro* tissues is the current challenge in the tissue engineering field [3, 4, 5].

The first stage to consider is the choice of the biomaterial or the bioink as a base for the tissue formation. This choice has an essential role in the whole process of determining the scaffold: printability, gelling, stability, mechanical properties, biocompatibility, biomimetics, bioactivity, and degradability or dissolution index [6, 7]. Biomaterials and bioinks are usually composed of soluble (bio)polymers with the capacity of crosslinking and forming scaffolds. The hydrogel’s highly hydrated environment (90 – 99% of water) shows a similar physical structure to the natural ECM allowing gases and liquids exchanges and, consequently, enabling cells to grow on and inside its meshes [8]. The most utilized bioinks in tissue engineering are the natural ones (collagen 26%, alginate 24%, hyaluronic acid 11%, and gelatin 10%) [8] as they are biocompatible and present biological characteristics, such as adhesion sites, which are important for modulating cell behavior [9]. However, natural hydrogels/polymers commonly present non-desirable mechanical properties as by definition they are highly swollen hydrophilic crosslinked materials. These non-desirable mechanical properties lead to a decay in their mechanical strength under cell culture conditions [10]. To address this issue, a blend of two or more biomaterials is usually needed for better results.

In this work, we used a combination of alginate and gelatin. Alginate is the second most used bioink due to the vast range of favorable characteristics for bioprinting and tissue engineering. It has shown satisfactory results on the delivery of small chemical drugs and proteins, cell culture, wound dressings, and tissue regeneration by delivering cells and proteins [11, 12, 13]. However, it presents poor mechanical properties and lacks cell bidding sites, so gelatin was added to address these issues. As shown by Giuseppe *et al*. (2018), varying the alginate and the gelatin concentrations tailors the hydrogel’s mechanical properties. Increasing the gelatin content in the alginate-gelatin hydrogel increases its elasticity [14, 15, 16]. Whereas increasing the alginate content increases its viscosity and stiffness [15, 16]. Besides, we can improve the biological characteristics of alginate by creating cell interactive alginate. Which can be produced by binding RGD peptides as side chains on alginate through carbodiimide chemistry [17]. The minimum amount of RGD necessary is cell type dependent. As reported by Alsberg *et al*. (2001) the transplantation of RGD-alginate containing primary rat calvarial osteoblasts into mice enhanced *in vivo* bone formation in comparison to alginate without RGD. Alginate is a potential biomaterial for bone tissue formation as it can deliver osteoinductive factors and/or bone-forming cells. Moreover, as a hydrogel, it has the capacity to fill irregular shapes, which is also an advantage in bone tissue engineering [12].

In the present study, we used the human osteosarcoma U2OS which is a well-established cell line having its proteome and genome characterized [19, 20]. Among 21 osteosarcoma cell lines, U2OS presented the highest cell growth rate and a high ability to migrate and invade [20]. Moreover, the high expression of E-cadherin allows this cell type to readily form spheroids as well as grow on mono or multilayers depending on the substrate [21, 22, 23]. Also, U2OS cells show the closest genome to the primary human osteoblast in comparison to other osteosarcoma cell lines [24]. Finally, even though U2OS has a mesenchymal origin, its flat fibroblast-like shape makes this cell type an important tool for cell imaging which is a crucial aspect when imaging cells on hydrogels [25, 26, 27]. All these characteristics made U2OS osteosarcoma the ideal model to study cell behavior and colonization of the fluid-phase tailored alginate-gelatin printings.

To date, freeform or embedded 3D printing allows the biofabrication of scaffolds from low viscosity hydrogels using a support bath. The fluid or fluid-like phase is usually composed of yield-stress fluids, sacrificial biomaterials, or granular liquid-like solids [28 - 32]. The fluid-phase 3D printing methodology we suggest in this work gives a step forward in the 3D (bio)fabrication field as we propose the utilization of the support bath containing an exchangeable cross-linking solution to customize hydrogels during the printing process. Tailoring the 3D printing is important to give each cell type the right stimuli to grow and differentiate. By printing the hydrogel within a fluid phase, a broad range of chemical and biological reactions can be done *in loco*. This new methodology allows the patterning of the scaffold biological, chemical, and mechanical parameters while printing by, for example, changing the crosslinker type and/or concentration, adding different molecules to the hydrogel in different concentrations to specific locations, programing incubation times, and doing all sort of chemical and biological reactions that needs an aqueous phase to happen. The objective of this work is to provide some examples on how to use the support bath to assist with the tailoring of the (bio)printings. Hence, the production of more elaborate biomaterials for *in vitro* studies which will enable a more accurate approach to basic research and drug screening assays.

## 2. MATERIALS AND METHODS

### 2.1 Materials

Alginic acid sodium salt type 1, Sodium Chloride, Calcium Chloride, and EDC (1-ethyl-3-(3-dimethylaminopropyl) carbodiimide hydrochloride) were purchased from Thermo Fisher Scientific; gelatin type B: bovine skin, fibronectin, and laminin were purchased from Sigma-Aldrich; MES buffer pH 7.0, Polyetherimide (PEI), and NHS (N-hydroxysuccinimide) were purchased from Alfa Aesar; MVG GRGDSP (RGD) was purchased from Novatech, RPMI 1640, fetal bovine serum (FBS), and trypsin/ EDTA were purchased from Gibco Life Technologies. U2OS osteosarcoma cell line (ATCC^®^ HTB-96^TM^) was used as bone-derived cell model.

### 2.2 Methods

#### Cell Culture and Seeding Protocol

U2OS cells were cultured in 75 cm^2^ flasks in complete RPMI 1640 supplemented with 10% FBS under a saturating humidified atmosphere at 37°C and 5% CO_2_. Subconfluent cultures were passaged at a ratio of 1:5 using 0.05% trypsin/ EDTA solution. High-density cells, 0.5 x 10^6^/cm^2^, were seeded onto the alginate-gelatin printings with or without ECM. Cells were also grown in spheroids formed on 0.8% agarose and the microcarriers: Cytodex^TM^ 1 (cross-linked dextran) (Cytiva) and macroporus Cytopore^TM^ 1 (cross-linked cotton cellulose) (Cytiva). Briefly, the microcarriers were first hydrated with PBS without Mg^2+^ and Ca^2+^ 5 ml/mg of microcarriers for 3 hours at room temperature. Then washed twice with same PBS 3 ml/mg of microcarriers, and sterilized in an autoclave at 115°C, 15 psi for 15 minutes. Prior to use the PBS was removed and the microcarriers were incubated with the same amount of RPMI 1640 medium + 10% FBS for 24 hours. 50 µl of the suspension was transferred to 12 well plate and 1 x 10^5^/cm^2^ of U2OS cells were seeded onto the microcarriers. Spheroids, Cytodex^TM^ 1, and Cytopore^TM^ 1 containing cells were kept in culture with complete RPMI 1640 supplemented with 10% FBS under a saturating humidified atmosphere at 37°C and 5% CO_2_ for 7 days. Then the spheroids, Cytodex^TM^ 1, and Cytopore^TM^ 1 containing cells were delivered onto the alginate-gelatin printings coated with RGD.

Cells growing on hydrogel substrates were monitored under an inverted phase contrast microscope (Olympus CKX53). Photomicrographs were taken by phase contrast microscopy using 4x, 10x, and 20x objective and images were analyzed by the software ImageJ/Fiji (NIH, USA).

#### Alginate-gelatin Hydrogel Preparation

Adapted from Alruwaili *et al* (2019), 2 or 7% gelatin was added to sterile 0.1 M MES buffer pH 7.0 + 0.3 M NaCl or complete RPMI 1640 + 10% FBS at 50°C and stirred for 10 minutes. Sodium alginate was then added at a final concentration of 6% and stirred thoroughly at 50°C until the hydrogel was homogeneous. Hydrogels were crosslinked overnight with 5 ml of 100 mM CaCl_2_ dissolved in deionized water.

#### Swelling and degradation Measurement

6% alginate + 2% gelatin in 0.1 M MES buffer + 0.3 M NaCl (H1), 6% alginate + 7% gelatin in 0.1 M MES buffer + 0.3 M NaCl (H2), 6% alginate + 2% gelatin in complete RPMI 1640 + 10% FBS (H3), and 6% alginate + 7% gelatin in complete RPMI 1640 + 10% FBS (H4) hydrogels molded into 10 mm diameter discs. The discs were weighed after molded (W_0_), after crosslinking with 100 mM CaCl_2_ (W_ACl_), and on day 1 (W_D1_), day 7 (W_D7_), day 14 (W_D14_), day 21 (W_D21_), and day 28 (W_D28_). The disks were kept under cell culture conditions incubated with RPMI 1640 supplemented with 10% FBS at 37°C and 5% CO_2_ with saturating humidity. Hydrogels were air-dried for 30 minutes before having their weight measured. Calculations were made of the percentage of W_0_ and experiments were done in triplicates and finished when 2/3 of the samples disintegrated.

#### 3D Printing, Printability Fidelity, and Printing Resolution Assays

The extrusion-based NAIAD 3D bioprinter, developed by the 3D Bioprinting Group from the Conway Institute at University College Dublin, was used to fabricate and tailor the alginate-gelatin hydrogel. Briefly, six-well plates were treated with 0.1% PEI overnight for hydrogel attachment purposes and washed 3 times with PBS to remove the excess PEI. Hydrogels were warmed at 37°C for 30 minutes in the bioprinter before the printing process. Alginate-gelatin printings were obtained using a 0.43 mm inner diameter (ID) nozzle gauge 23 (Weller KDS2312P) under 3 bars of pressure and 1mm/s of speed in air or fluid-phase containing 100 mM CaCl_2_.

For the 2D printability assay, 6% alginate + 2% gelatin in 0.1 M MES buffer pH 7.0 + 0.3 M NaCl, hydrogel H1, was printed in a square wave shape using a 0.43 mm ID nozzle gauge 23 under 1-6 bars of pressure and 1-5 mm/s of speed in air or fluid-phase containing 5 ml of 100 mM CaCl_2_. The printings had the linearity of the strands and the internal angle of the corners analyzed qualitatively, where: red= designed printing not obtained, yellow: low fidelity of lines and corners, green= good fidelity of lines and low fidelity of corners, dark green= good fidelity of lines and corners. For the quantification of the printing resolution, we measured the spread of the printing strand. The lines were printed using the same nozzle under 4 bars of pressure and 1-5 mm/s of speed in air or fluid-phase containing 5 ml of 100 mM CaCl_2_ and had its diameter measured immediately after printing. Data was analyzed by the software ImageJ/Fiji (NIH, USA).

For the 3D printability assay, hydrogel H1 was printed in 3 layers of wall thickness and 15 layers of height using the same 0.43 mm ID nozzle gauge 23 under 3 bars of pressure and 1 mm/s of speed in air or fluid-phase containing 7 ml of 100 mM CaCl_2_. Photomicrographs were taken on a modular Stereo Microscope Zeiss SteREO Discovery V20.

#### Alginate-gelatin Hydrogel Coating with ECM

Adapted from Rowley *et al* (1999), alginate-gelatin hydrogel printings were incubated with either RGD (0, 60, 120, and 240 µM/ml) or fibronectin (1 µg/ml) or laminin (1 µg/ml) + 5% EDC + 2.5% NHS diluted in either 20 or 100 mM CaCl_2_ for 1 hour. To assess the minimum incubation time was necessary for an efficient incorporation of RGD into alginate via carbodiimides chemistry, alginate-gelatin hydrogel was molded into 10 mm diameter discs and incubated with 120 µM/ml of RGD + 5% EDC + 2.5% NHS diluted in 100 mM CaCl_2_ for 0, 15, 30, 60 minutes and overnight. Cell attachment was measured by the percentage of the reduction of alamarBlue^TM^ (Invitrogen) assay on day 3 of cell culture. The hydrogel without RGD was used as a control for calculations purposes. Calculations were given as a percentage of control. Hoechst 33342 (Sigma Aldrich) staining was performed on day 7 of cell culture after same incubations time with RGD + carbodiimides in a cross-linkable bath. Photomicrographs were taken on the Olympus CKX41 inverted microscope with U-RFLT50 power supply unit using the 10x objective.

#### Hydrogel Patterning

The NAIAD 3D bioprinter was used to print the 6% alginate + 2% gelatin in 0.1 M MES buffer pH 7.0 + 0.3 M NaCl, hydrogel H1, using a 0.43 mm ID nozzle gauge 23 under 3 bars of pressure and 1 mm/s of speed in fluid-phase containing CaCl_2_. Before printing, the cell culture plates were incubated overnight with PBS + 0.1% of PEI and prior to print, the wells were washed 3 times with PBS to remove the excess PEI and the cross-linkable solution was added to the wells.

To test the effect of RGD on U2OS cells, a one-layer flat surface was half printed under 5 ml of 100 mM CaCl_2_ added with 120 µM/ml of RGD + 5% EDC + 2.5% NHS and incubated for 1 hour for RGD incorporation into alginate. After that, the half printing was washed 3 times with PBS to remove the non-attached RGD and the second half of the printing was printed under 5 ml of 100 mM CaCl_2_ without RGD. After crosslinking overnight, the hydrogel was washed 3 times with PBS and U2OS cells were seeded (Fig. 1A).

**Figure 1:**
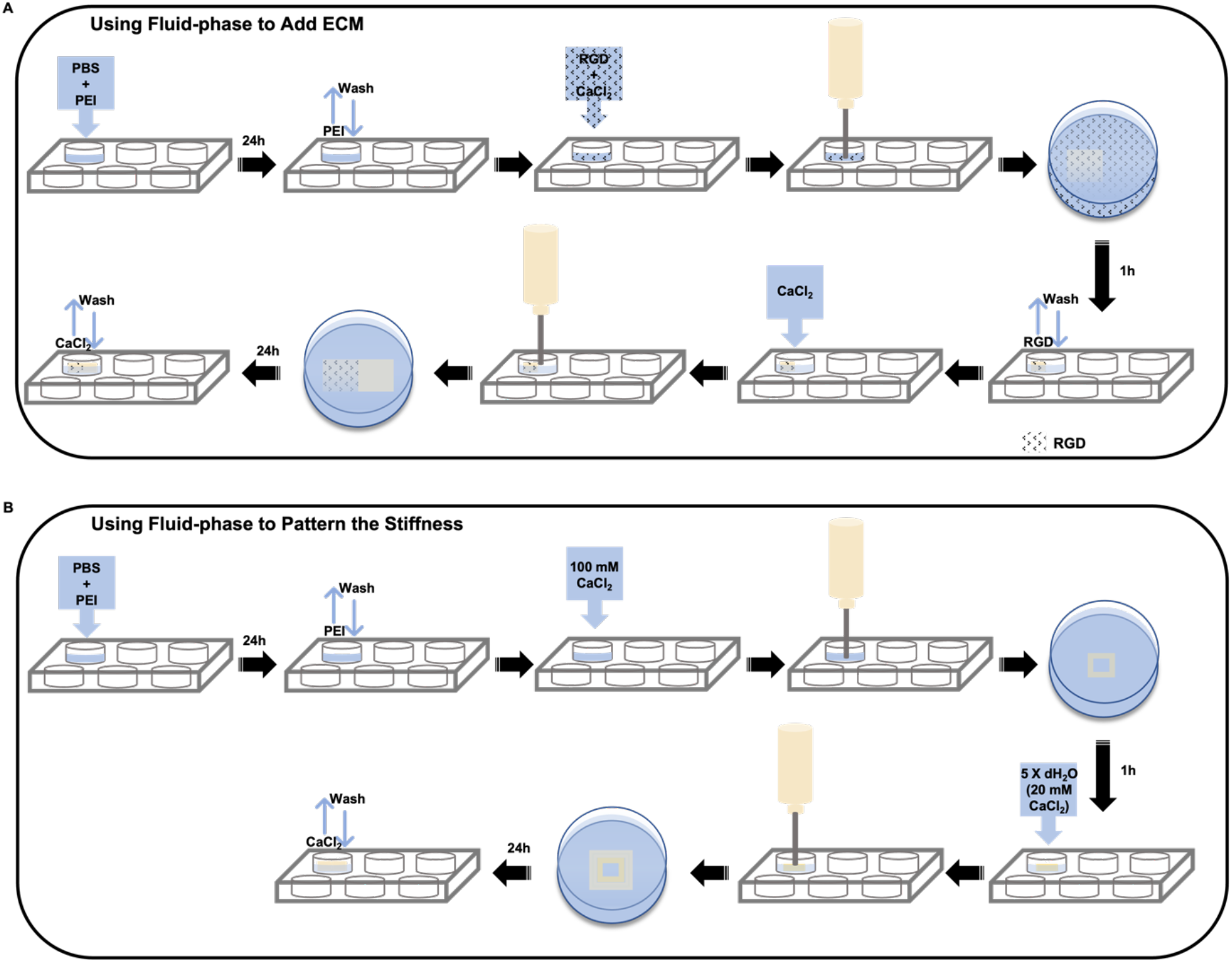
Fluid-phase methodology scheme. **(A)** A model of biological patterning of alginate-gelatin hydrogel with RGD by fluid-phase. Plates were incubated overnight with PBS + 0.1% of PEI, the wells were washed 3 times with PBS to remove the excess PEI and 100 mM CaCl_2_ + 120 µM RGD/mL + carbodiimides was added. Half of the printing was obtained at 37°C and incubated for 1 hour for RGD incorporation into alginate. Posteriorly, the printing was washed 3 times with PBS to remove the RGD not attached, and the second half of the printing was obtained under 100 mM CaCl_2_ without RGD. After overnight crosslinking the printing were washed and the cells were seeded. **(B)** A model of physical patterning of alginate-gelatin hydrogel varying the crosslinker concentration with fluid-phase. Plates were incubated overnight with PBS + 0.01% of PEI, the wells were washed 3 times with PBS to remove the excess PEI and 100 mM CaCl_2_ was added. Half of the printing was obtained and incubated for 1 hour to cross-link. Posteriorly, the crosslinker solution was diluted 5X and the second half of the printing was obtained under 20 mM CaCl_2_. After crosslinking the printings were washed and the cells were seeded.

To test the effect of stiffness on U2OS cells, a one-layer open square was printed under 2 ml of 100 mM CaCl_2_ and incubated for 1 hour for crosslinking, then the cross-link solution was diluted to 20 mM, and a second outer one-layer open square was printed and incubated for 1 hour for crosslinking. Next, the printing was washed 3 times with PBS and U2OS cells were seeded (Fig. 1B). RGD + carbodiimides concentrations were kept constant at 120 µM/ml of RGD + 5% EDC + 2.5% NHS during the whole printing and crosslinking duration.

#### Cell Viability Assay

LDH Cytotoxicity Detection Kit plus (Roche Diagnostics) was used to quantify U2OS cell death on day 1 and day 7 of cell culture. Briefly, cells were incubated for 24 hours with RPMI 1640 supplemented with 1% FBS, centrifuged, and the amount of LDH present on the cell culture medium was measured at 490 nm by spectrophotometer (SpectraMax M3). Positive and negative controls were done on conventional 2D cell culture on plastic plates. Positive control was treated with 2% Triton X-100 (Sigma Aldrich). Calculations were given as a percentage of control.

#### Cell Quantification

U2OS cells growing on alginate-gelatin hydrogels were washed twice with PBS to remove non-attached cells and cell culture medium residue. The hydrogels containing cells were treated with 0.05% Trypsin/EDTA for 5 mins and cells were counted in a Neubauer chamber.

#### AlamarBlue^TM^ Assay

A standard protocol was used for the alamarBlue^TM^ assay. In brief, 10% of alamarBlue^TM^cell viability reagent (Invitrogen) was added to the complete RPMI 1640 +10% FBS medium on day 3 or 7 of cell culture and incubated for 4 hours. Next, the cell culture medium containing alamarBlue^TM^ was centrifuged and 100 µl transferred to a 96 well plate. The absorbance of the reduced alamarBlue^TM^ was measured at 570 nm and 600 nm by spectrophotometer (SpectraMax M3) and the cell metabolic activity was calculated according to the manufacture’s instruction. For calculations purposes, cells growing on hydrogels crosslinked with 100 mM CaCl_2_ without RGD were used as control. Calculations were given as a percentage of control.

#### Hoechst 33342 and PI Double Staining (live staining)

U2OS cells were incubated with 4 µM Hoechst 33342 (Sigma Aldrich) and 1 µg/ml PI (Sigma Aldrich) for 15 minutes and images were taken with the Olympus CKX41 inverted microscope with U-RFLT50 power supply unit using the 10x objective. Cells/field was measured by the intensity of fluorescence quantified by ImageJ/Fiji (NIH, USA) for live dead assay. Calculations were given as a percentage of control.

#### Rheology Test

Hydrogel H1, 6% alginate + 2% gelatin in 0.1 M MES buffer pH 7.0 + 0.3 M NaCl, was molded into 10 mm diameter discs with ∼ 5 mm height and crosslinked overnight with 5 ml of either 20, 100 or 300 mM CaCl_2_. Prior to the tests, the crosslinked disks were washed with PBS 3 times. The amplitude sweep test was performed on MCR301 rheometer (Anton Paar). All the measurements were done using a parallel plate with a 25 mm diameter top plate geometry with gap varying between 4 - 6 mm at 37°C. The amplitude sweep was performed at a strain ranging from 0.01 to 100% and a frequency of 1 Hz. Storage (G’) and loss (G’’) moduli of the hydrogels were measured to determine the linear viscoelasticity region (LVR). Prior to the measurements, hydrogels were equilibrated at 37 °C for 2 minutes.

#### Statistical analysis

All statistical data processing was performed using multiple comparison tests on either Student’s *t*-test or One-way ANOVA or Two-way ANOVA on GraphPad Prism 9 software. Differences between the groups were considered reliable if p< 0,05 and values were expressed as mean ± SD of at least 3 independent experiments.

## 3. RESULTS

To determine which alginate-gelatin hydrogel substrate would give support to growing bone tissue *in vitro*, we tested 4 different formulations for swelling and degradation during 28 days under cell culture conditions (Table 1). Cell differentiation into osteoblasts and beginning of mineralization occur in general between 14 and 21 days of cell culture for human primary mesenchymal cells and continuous bone-derived cells [36 - 39]. Prior to incubation with cell culture medium, all hydrogels were molded into 10 mm discs and weighed before and after overnight crosslinking with 100 mM CaCl_2_ (W_0_ and W_ACl_, respectively). The hydrogel H1, prepared with 6% alginate + 2% gelatin in 0.1 M MES buffer + 0.3 M NaCl, was the only formulation that kept its integrity for 21 days presenting 70% of swelling on day 21 (Fig. 2A). The hydrogels prepared with higher gelatin ratio (H2 and H4) presented weaker mechanical strength and degraded faster than their counterparts prepared with lower gelatin ratio (H1 and H3, respectively). Increasing the gelatin content alters the hydrogel mechanical properties as gelatin behaves as a liquid at 37°C. This leads to a decrease in hydrogel strength and, consequently, an increase in the degradability/dissolution of those substrates. The solvent used to prepare the hydrogels also played an important role as the formulations prepared with the complete cell culture medium RPMI 1640 + 10% FBS (H3 and H4) lost strength faster than their counterparts prepared with 0.1 M MES buffer + 0.3 M NaCl (H1 and H2, respectively) (Fig. 2A). To study cell colonization of the alginate-gelatin substrates, we added RGD to the hydrogels as alginate lacks cell binding sites and according to Alsberg *et al*. (2001) osteoblasts show an increase on cell growth in the presence of RGD peptides. To determine if a higher gelatin content would provide cells with even more variety of cell binding sites and increase cell attachment and growth, we quantified the amount of U2OS cells growing on the hydrogels H1 (2% gelatin) and H2 (7% gelatin) on day 7 of cell culture. Our results showed that increasing the gelatin content in the alginate hydrogel was not beneficial neither for the hydrogel stability nor for cell growth as hydrogel H1 presented 2.83-fold greater cell growth in comparison to H2 (Fig. 2B). Therefore, hydrogel H1 showed to be a more suitable substrate to keep bone tissue developing *in vitro* over time among the formulations tested in this study. Next, 2D and 3D printings were created to observe cell colonization of the alginate-gelatin hydrogel H1. U2OS cells were kept in culture for up to 20 days and showed good attachment and growth on the printings. A substantial cell growth was seen on both large 2D printings and narrow 3D printings (Fig. 2C). However, fine structures do not seem to last long periods in culture. Whereas larger structures showed good stability over time and allowed cells to increase colonization and grow in multilayers and aggregates in a time-dependent manner (Fig. 2C). Growing in multilayers is the first and an important step for bone differentiation and mineralization [38]. Finally, the cells showed around 13% of death rate on day 20 quantified by the relative fluorescent intensity of cells stained with PI against the total cell number stained with Hoechst 33342 (Fig. 2D).

**Table 1:** List of the alginate-gelatin hydrogels tested in this work with the respective description of their formulations.

| **HYDROGEL** | **FORMULATION** |
| --- | --- |
| **H1** | 6% alginate + 2% gelatin in 0.1 M MES + 0.3 M NaCl |
| **H2** | 6% alginate + 7% gelatin in 0.1 M MES + 0.3 M NaCl |
| **H3** | 6% alginate + 2% gelatin in complete RPMI medium + 10% FBS |
| **H4** | 6% alginate + 7% gelatin in complete RPMI medium + 10% FBS |

**Figure 2:**
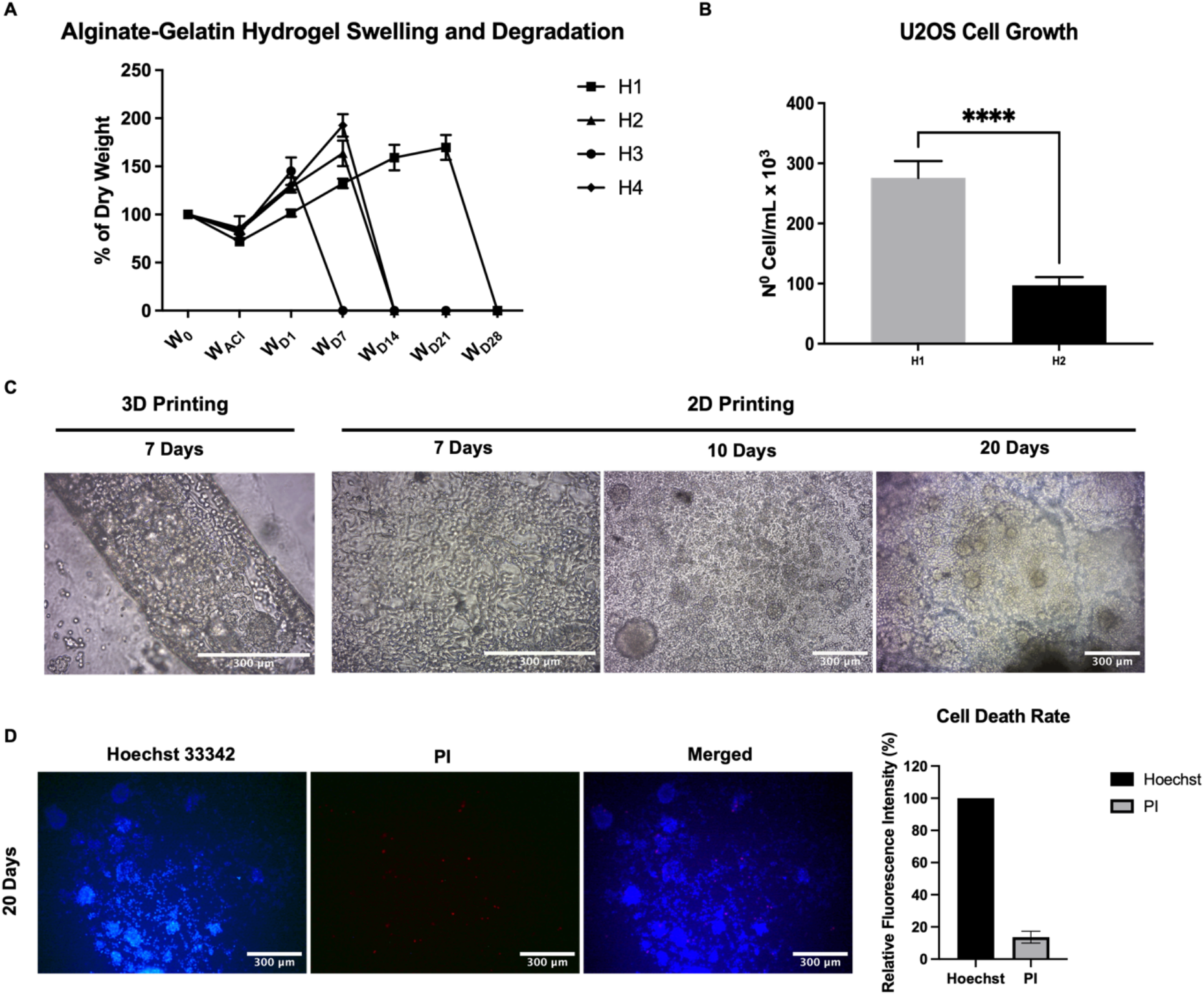
(A) (H1) 6% alginate + 2% gelatin in 0.1 M MES buffer + 0.3 M NaCl, (H2) 6% alginate + 7% gelatin in 0.1 M MES buffer + 0.3 M NaCl, (H3) 6% alginate + 2% gelatin in complete RPMI 1640 + 10% FBS, and (H4) 6% alginate + 7% gelatin in complete RPMI 1640 + 10% FBS. Hydrogels were moulded into 10 mm diameter discs and weighted (W_0_= 100%) and air-dried and weighed after crosslinking with 100 mM CaCl_2_ (W_ACl_), day 1 (W_D1_), day 7 (W_D7_), day 14 (W_D14_), day 21 (W_D21_), and day 28 (W_D28_). The samples were kept under cell culture conditions incubated with RPMI 1640 supplemented with 10% FBS at 37°C and 5% CO_2_ with saturating humidity (n= 6, graph of mean ±SD). **(B)** Quantification of U2OS cells attached to 6% alginate + 2% gelatin (H1) and to 6% alginate + 7% gelatin (H2) hydrogels on day 7 of cell culture. Cells were counted in a Neubauer chamber after treatment with trypsin (n= 9, t test, p****<0.0001). **(C)** Phase contrast images of U2OS cells on the hydrogel H1 added with 120 µM RGD/mL and crosslinked with 100 mM CaCl_2_. Pictures were acquired on day 7, 10, and 20 of cell culture. Day 7: magnification of 20x; day 10 and 20: magnification of 10x. **(D)** Live Dead assay done by Hoechst /PI double staining of U2OS cells on the hydrogel H1 added with 120 µM RGD/mL on day 20 of cell culture and quantification. The quantification was done with the percentage of relative fluorescence intensity by the ImageJ/Fiji software (n= 5, t test, p****< 0.0001).

To optimize cell growth and colonization of hydrogel H1, the extrusion-based 3D bioprinter NAIAD was used to acquire 2D and 3D printings in fluid-phase. The benefits of printing in fluid-phase are that we can do a vast range of biological and chemical reactions during the printing process *in loco*. Hence patterning the substrate to improve and guide cell colonization and behavior onto the printings. In this context, first, to investigate the printability of the hydrogel H1 in air and in fluid-phase, we used the crosslinker CaCl_2_ (100 mM) as aqueous support bath to obtain the 2D printings and the same solution added with 0.1% PEI to obtain the 3D printings (Supplementary Video 1). PEI increases the attachment between layers during the construction of the scaffolds. Printability was first analyzed qualitatively and then by the spread of the strand printed in air and fluid-phase. Printing in fluid-phase, containing the crosslinker, enlarges the spectrum of the printing settings mainly within lower pressure (pressure vs. speed) in comparison to printing in air (Fig. 3A). However, high speeds decrease the attachment between the hydrogel and the plate or layers when printing in fluid-phase which also leads to a decrease in printing fidelity. Overall, the fluid-phase increased the printing fidelity by keeping the lines, corners, layers, and lumens in 2D and 3D printings more defined than the air printing under the same conditions (Fig. 3B); and finally, the fluid-phase increased printing resolution by limiting the spreading of the printed lines as it cross-links the hydrogel immediately after printing. Therefore, in general, fluid-phase keeps the printings more faithful to the designed structures and would be especially valuable for printing soft hydrogels that collapse easier than the stiffer ones (Fig. 3C).

**Figure 3:**
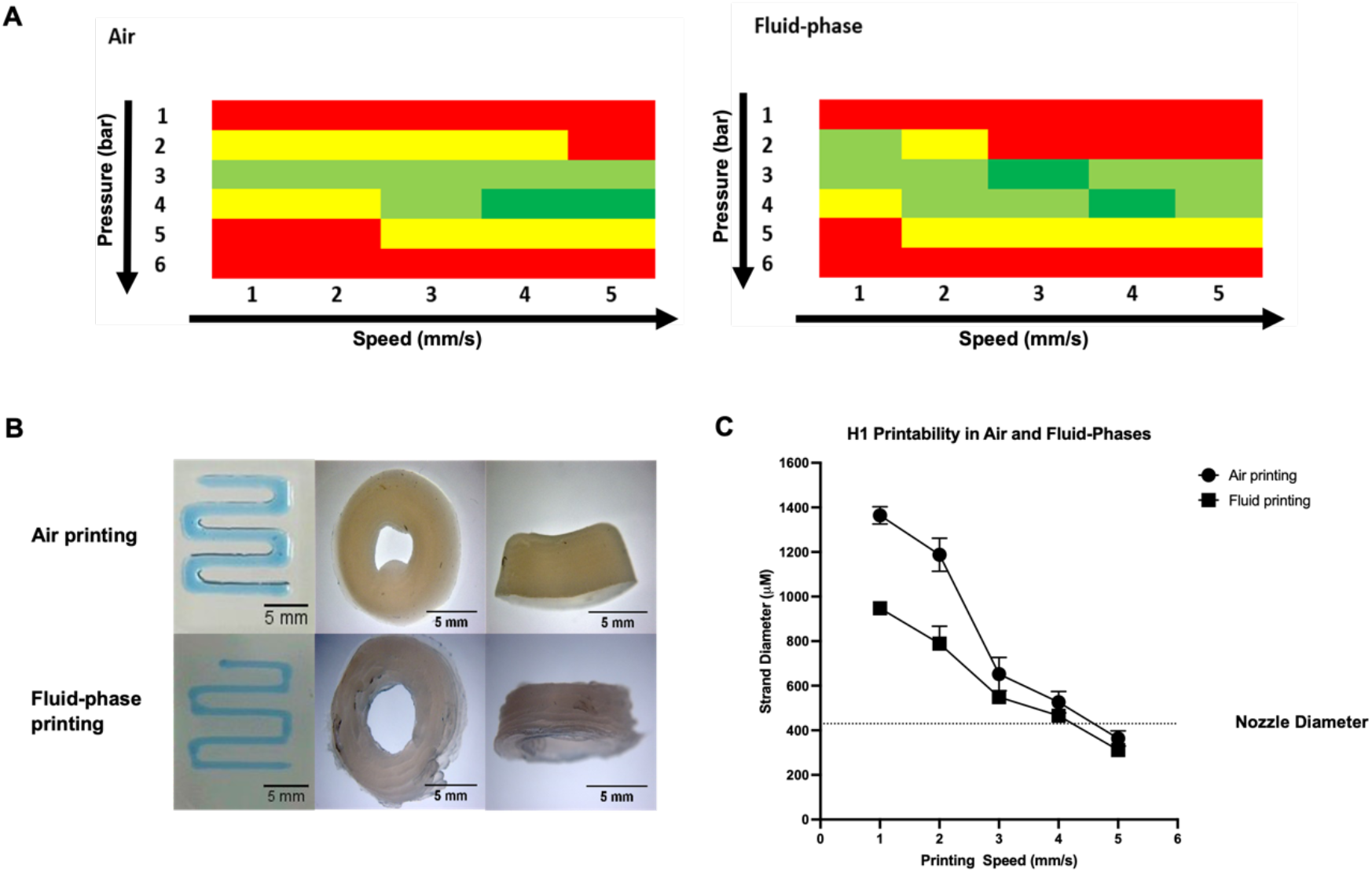
(A) 6% alginate + 2% gelatin, hydrogel H1, printability in air and fluid-phase under 1-6 bars of pressure and 1-5 mm/s of speed. Qualitative printing fidelity chart. Red= designed printing not obtained, yellow: low fidelity of lines and corners, green= good fidelity of lines and low fidelity of corners, dark green= good fidelity of lines and corners. **(B)** 6% alginate + 2% gelatin, hydrogel H1, 2D printing in air and fluid-phase under 3 bars and 3 mm/s (square wave) and 3D printing in air and fluid-phase under 3 bars and 1 mm/s (15 layers high and 3 layers thick tube). **(C)** 6% alginate + 2% gelatin, hydrogel H1, strand diameter in air and fluid-phase under 4 bars of pressure and 1, 2, 3, 4, and 5 mm/s of speed (n= 7, Two-way ANOVA, 1 and 2 mm/s: p****< 0.0001, 3: p*< 0.05, 4 and 5: not significant).

As aforementioned, an important factor for growing U2OS cells on alginate hydrogel substrates is the presence of cell binding sites on the printed scaffolds. To assess the effect of ECM on U2OS cell growth, the alginate-gelatin hydrogel H1 was coated with 3 different molecules: RGD, fibronectin, and laminin. All 3 molecules boosted U2OS cell growth and increased spread on the hydrogels’ surfaces. Moreover, no significant difference was observed between them (Fig. 4A). Therefore, we continued the study using RGD to pattern our printings by fluid-phase. However, it was first necessary to determine the incubation time was needed to efficiently incorporate RGD molecules into alginate through carbodiimide chemistry in a cross-linkable aqueous support bath. For this purpose, H1 printings were incubated with RGD + EDC + NHS in 100 mM CaCl_2_ for different times (0, 15 min, 30 min, 60 min, and overnight). The number of cells attached to each substrate was assessed by the percentage of alamarBlue^TM^ reduction on day 3 of cell culture (Fig. 4B) and the Hoechst 33342 staining on day 7 of cell culture (Fig. 4C). As a result, we observed that a minimum of 1 hour of incubation with RGD + carbodiimides is necessary for the optimization of the reaction in a cross-linkable fluid-phase. In addition, by coating the H1 hydrogel printings with RGD we can deliver U2OS cells through different means onto the substrates. In this study, we tested the delivery of cells by spheroids, Cytodex^TM^ 1, and Cytopore^TM^ 1, besides conventional cell seeding protocol. We observed that from day 1 cells bonded to the hydrogel and from day 2 on, they started colonizing the surface of our substrates. Cells massively migrated from the delivering structures to the hydrogel surface within 7 in cell culture (Fig. 4D).

**Figure 4:**
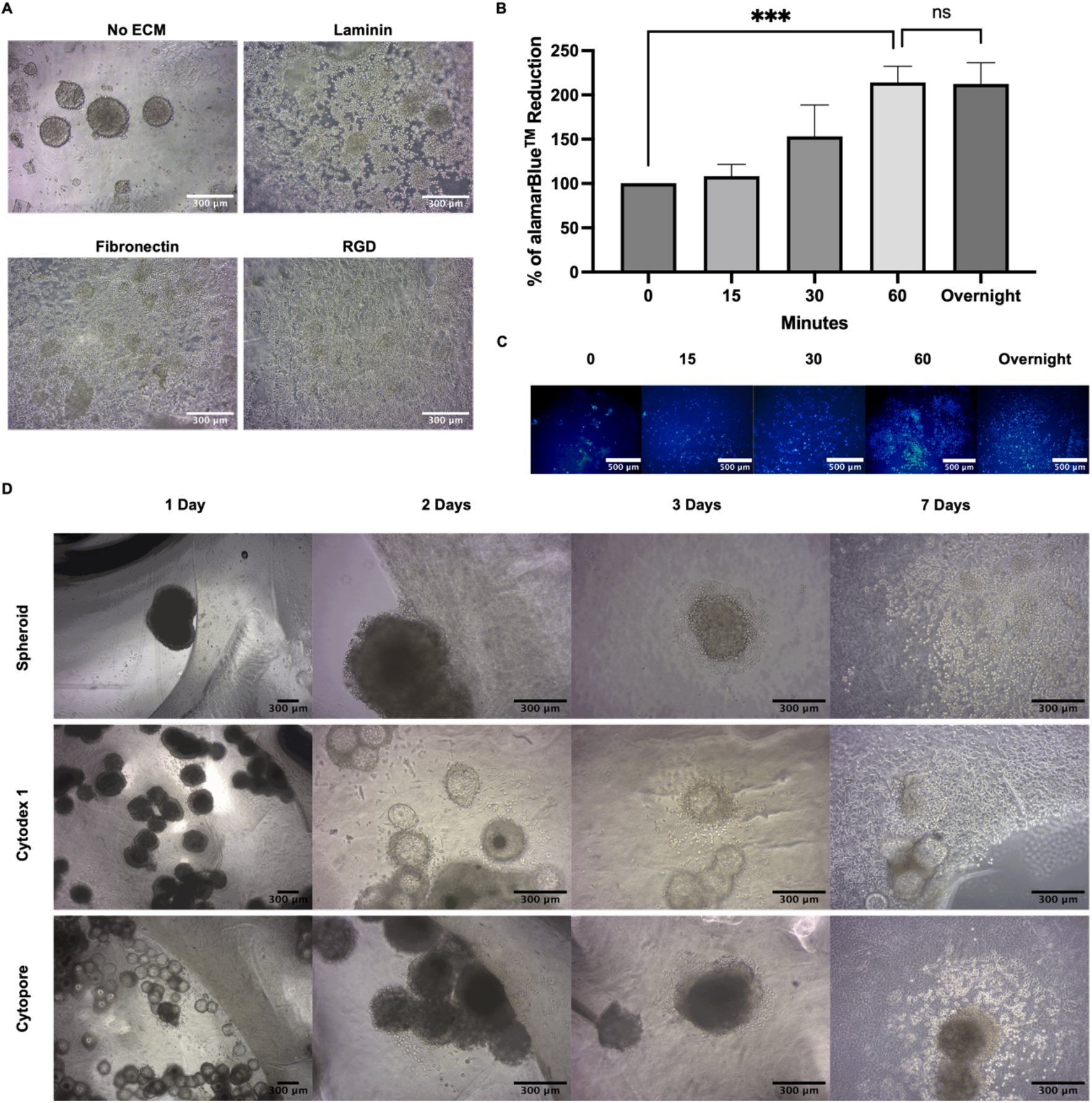
(A) U2OS cells on 6% alginate + 2% gelatin, hydrogel H1, crosslinked with 100 mM CaCl_2_ and coated with either 120 µM of RGD/mL or 1 µg of fibronectin/mL or 1 µg of laminin/mL, or without ECM. Images were acquired on day 7 of cell culture. Magnification of 10x. **(B)** Quantification of the percentage of alamarBlue^TM^ reduction of U2OS cells growing on the 6% alginate + 2% gelatin, hydrogel H1, on day 3 of cell culture. Hydrogels were incubated for 0, 15, 30, and 60 minutes, and overnight with 120 µM of RGD/ml + carbodiimides in 100 mM CaCl_2_. Calculations were done as percentage of the control (hydrogel without RGD was considered 100%) (n= 3, One-way ANOVA, p****< 0.001). **(C)** Hoechst staining of U2OS cells on 6% alginate + 2% gelatin, hydrogel H1, after 0, 15, 30, and 60 minutes, and overnight incubations with 120 µM of RGD/ml + carbodiimides in 100 mM CaCl_2_. Images were acquired on day 7 of cell culture. Magnification of 10x. **(D)** U2OS cells delivered to the hydrogel H1printings by spheroids, Cytodex^TM^ 1 or Cytopore^TM^ 1. First row: U2OS spheroid seeded onto 6% alginate + 2% gelatin, hydrogel H1, coated with 120 µM of RGD/mL. Second row: Cytodex^TM^ 1 containing U2OS cells seeded onto 6% alginate + 2% gelatin, hydrogel H1, coated with 120 µM of RGD/ml. Third row: Cytopore^TM^ 1containing U2OS cells seeded onto 6% alginate + 2% gelatin, hydrogel H1, coated with 120 µM of RGD/ml. Images were acquired on days 1, 2, 3, and 7 of cell culture. Images of day 1 were acquired with 4X and images of days 2, 3, and 7 were acquired with 10x objective.

To guide cell colonization on our substrates, the NAIAD 3D bioprinter, combined with the fluid-phase methodology, was used to add RGD molecules to specific parts of the hydrogels during the printing process. Our methodology for biological patterning of alginate-gelatin hydrogels with ECM was shown to be efficient to compartmentalize the U2OS growth to those parts containing RGD. In addition, cell death, shown by the PI staining, was not significant neither on the parts coated with RGD nor on parts without RGD (Fig. 5A). The coating with RGD increased 2.97 times the U2OS cell growth on the alginate-gelatin hydrogel H1 (Fig. 5B).

**Figure 5:**
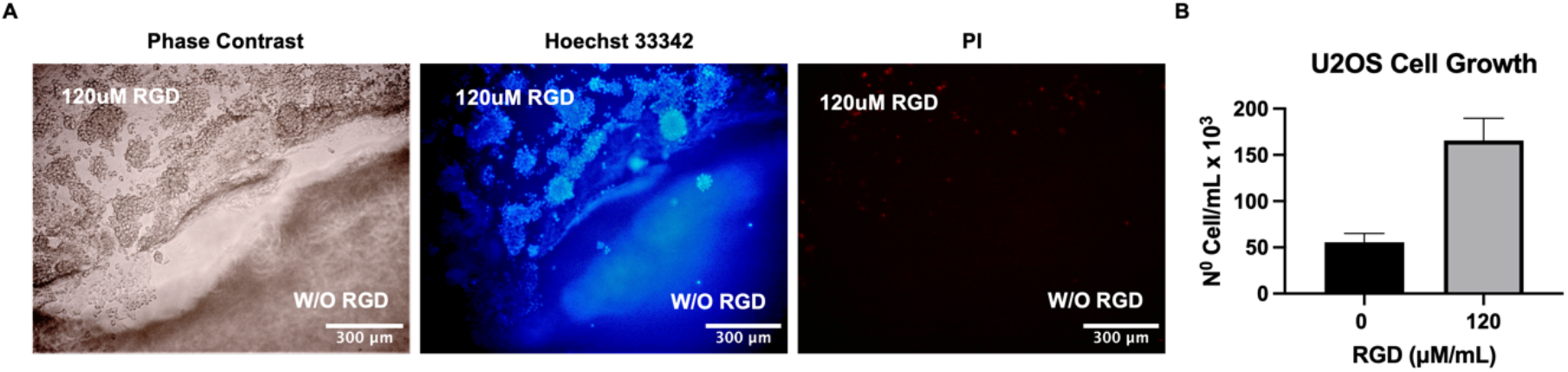
(A) Light microscopy (phase contrast) and Hoechst 33342/ PI double staining of U2OS cells on 6% alginate + 2% gelatin, hydrogel H1, tailored by fluid-phase. Hydrogels were coated on pre-determined areas with 120µM of RGD/mL. Images were acquired on day 5 of cell culture. Magnification of 10x. **(B)** Quantification of U2OS cells attached to the alginate-gelain hydrogel H1 coated with 120µM of RGD/mL and without RGD on day 7 of cell culture. Cells were counted in a Neubauer chamber after treatment with trypsin (n= 8, Student t test, p****<0.0001).

As previously reported by Jabbari *et al.* (2015), U2OS cells need a stiff hydrogel (∼50 KPa) to optimize cell growth. Therefore, the NAIAD 3D bioprinter, combined with the fluid-phase methodology, was also used to tailor the stiffness of the alginate-gelatin hydrogel H1. This methodology also proved to be efficient at compartmentalizing cell growth to pre- determined spots of the printings. By printing a stiffer substrate under a more concentrated crosslinking bath and a softer substrate under a less concentrated crosslinking bath, we could guide the growth of U2OS cells to stiffer spots of the printings (Fig. 6A). Corroborating the previous data, the stiffer substrate showed an increase of 3.26 folds on the total number of cells and an increase of 2 folds on the cell metabolic activity on day 7 of cell culture in comparison to the softer substrate (Fig. 6B, C). This increase in the cell number on the stiffer substrate was observed even though the hydrogel crosslinked with 100 mM CaCl_2_ caused an increase in cell death of 2.39 folds in comparison to 20 mM CaCl_2_ on day 1. On day 7 of cell culture, both substrates presented a similar cell death rate of around 5% (Fig. 6D).

**Figure 6:**
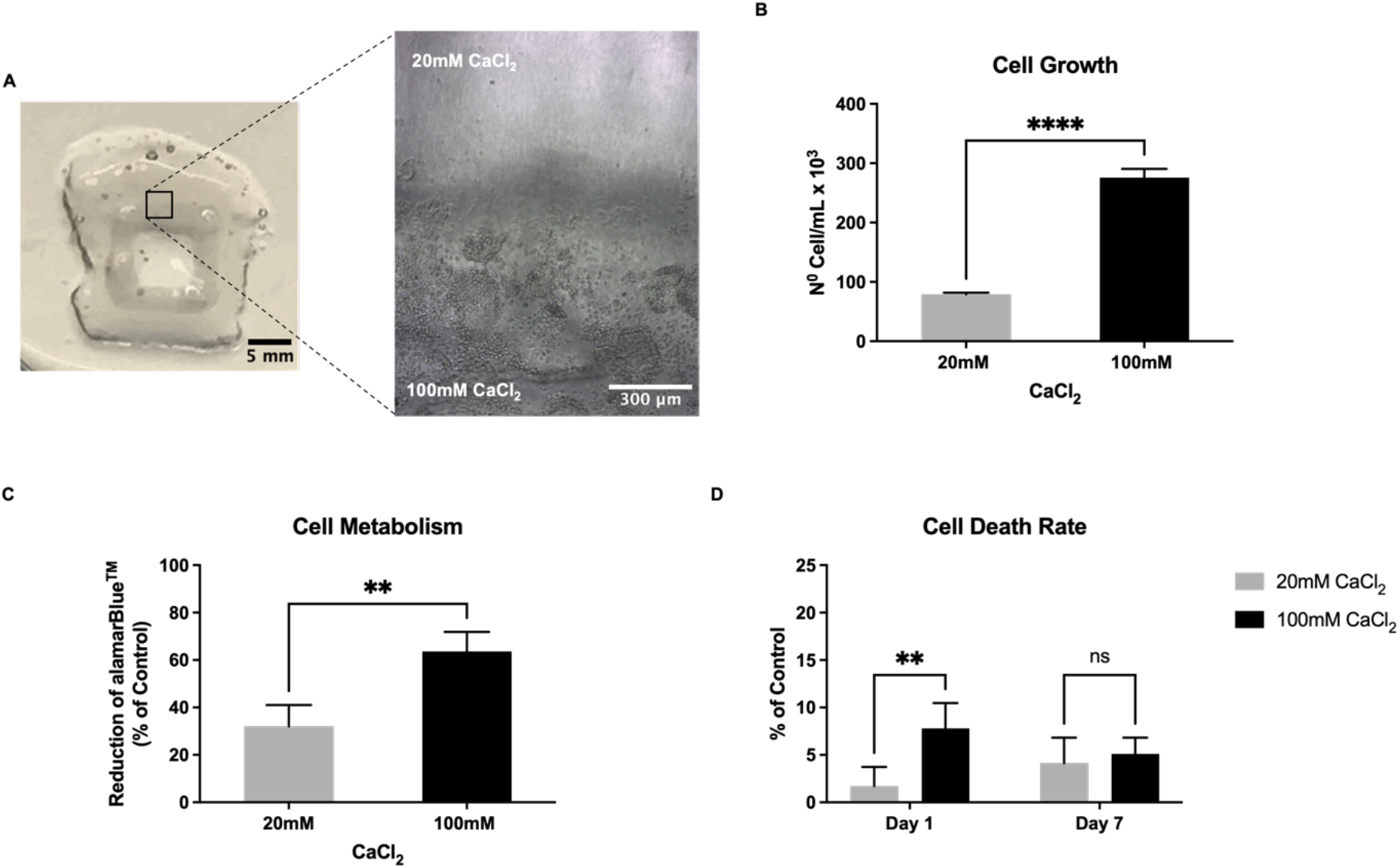
(A) One-layered open square printing showing its stiffness tailored by fluid-phase. Inner square printed under 100 mM CaCl_2,_ and outer square printed under 20 mM CaCl_2_. The whole printing was coated with 120µM of RGD/ml. U2OS cells were seeded onto the tailored 6% alginate + 2% gelatin, hydrogel H1, and images were acquired on day 5 of cell culture. Magnification of 10x. **(B)** Quantification of U2OS cells on the hydrogel H1 crosslinked with either 20 mM or 100 mM CaCl_2_ on day 7 of cell culture. Hydrogels were coated with 120µM of RGD/ml. Cells were counted in a Neubauer chamber after treatment with trypsin (n= 8, Student t test, p****<0.0001). **(C)** Quantification of the % of alamarBlue^TM^ reduction of U2OS cells on the hydrogel H1 crosslinked with either 20 mM or 100 mM CaCl_2_ on day 7 of cell culture. Hydrogels were coated with 120µM of RGD/ml (n= 4, Student t test, p**< 0.01). **(D)** LDH Cytotoxicity Assay of U2OS cells on the hydrogel H1 crosslinked with either 20 mM or 100 mM CaCl_2_ both coated with 120µM of RGD/ml on day 1 and 7 of cell culture (n= 5, Student t test, day 1 p**<0.01, day 7 not significant).

Next, to access the rheological aspects of the hydrogel H1 crosslinked with increasing concentrations of CaCl_2_, discs of the hydrogel were moulded into 10 mm discs and crosslinked with 20, 100 or 300 mM CaCl_2_. Printings obtained in fluid-phase presented roughness on their surfaces unsuitable for accurate rheological tests (Fig. 7A). Moreover, printing in fluid-phase containing 300 mM CaCl_2_ showed an important lack in printing fidelity. The rheological results showed that the crosslinking with 300 mM CaCl_2_ resulted in a Young’s modulus of 114.33 ± 1.53 KPa while the crosslinking with 100 mM CaCl_2_ resulted in a Young’s modulus of 58.83 ± 4.51 KPa, and, finally, the crosslinking with 20 mM CaCl_2_ resulted in a Young’s modulus of 4.35 ± 2.14 KPa, which is considered to be too soft for growing U2OS cells [39] corroborating our previous findings (Fig. 7B, C). Besides, we could confirm that the crosslinker concentration can modify the strength and elasticity of the hydrogel. The crosslinking with 300 mM CaCl_2_ showed the highest G’ LVR and the biggest difference between storage and Loss moduli in comparison to the crosslinking with 100 and 20 mM CaCl_2_ (Fig. 7D – F). Changing crosslinker concentrations showed to be an efficient and cost-less way to create different stiffness and improve mechanical properties of alginate-gelatin hydrogel.

**Figure 7:**
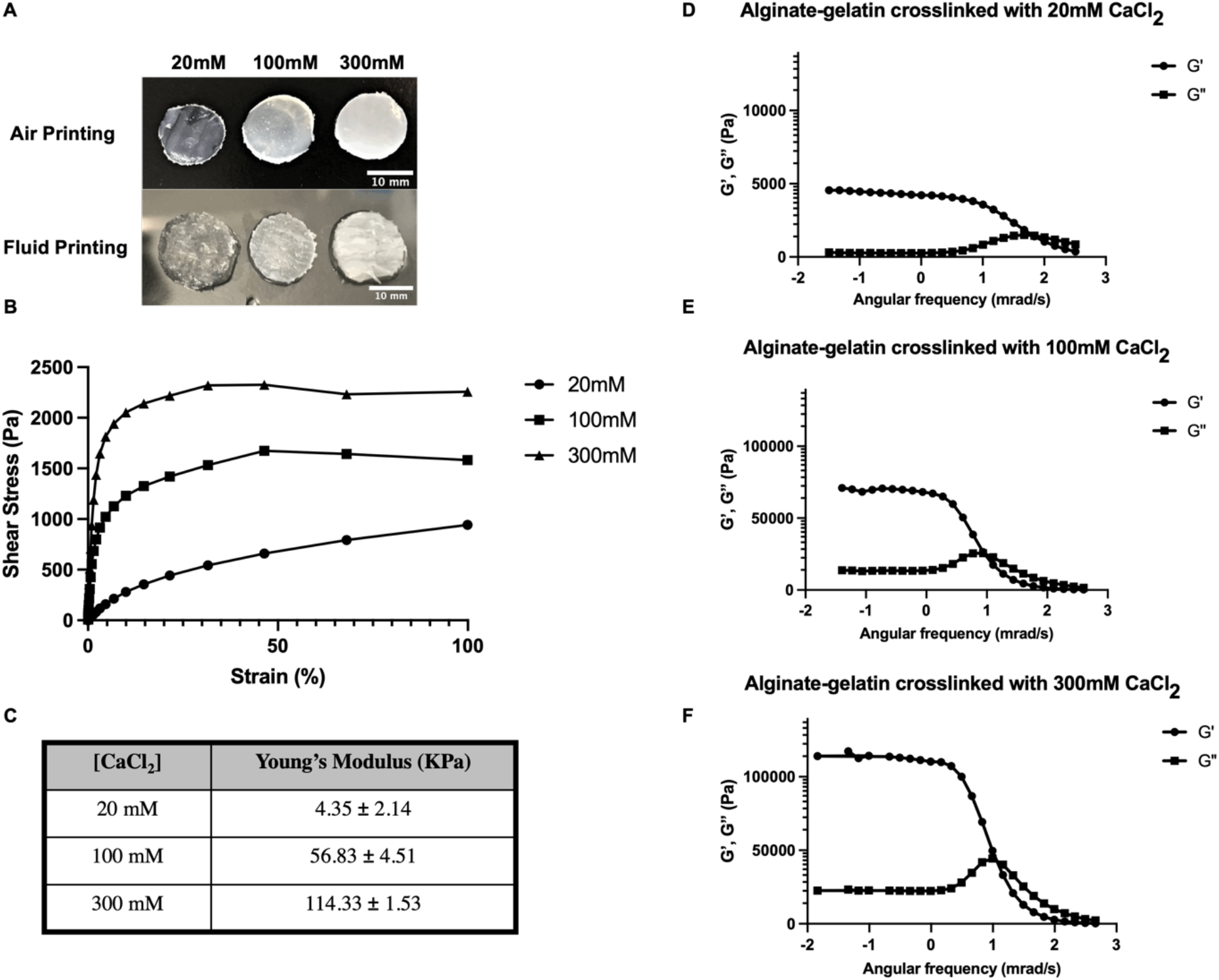
Rheological tests of the 6% alginate + 2% gelatin, hydrogel H1, crosslinked with 20, 100 or 300 mM CaCl_2_. **(A)** Hydrogel H1 printed in air and fluid-phase and crosslinked with 20, 100 or 300 mM CaCl_2_. **(B)** Shear stress vs strain (%) of the hydrogel H1 moulded into 10 mm disks and crosslinked with 20, 100 or 300 mM CaCl_2_. **(C)** Young’s modulus table of the hydrogel H1 crosslinked with 20, 100 or 300 mM CaCl_2_. **(D, E, F)** Storage (G’) and loss (G”) moduli of the hydrogel H1 crosslinked with 20 mM (E), 100 mM (D) or 300 mM (F) CaCl_2_ plotted against the angular frequency.

Finally, by combining increasing stiffness and increasing RGD content on the alginate-gelatin hydrogel H1, we could improve the cell metabolic activity by up to 3.17 folds. Depending on the stiffness of the substrate, we can see a different response to the RGD concentration on the cell metabolism. On the soft hydrogel (4.35 KPa), crosslinked with 20 mM CaCl_2_, cells do not present high cell growth rate, which is shown by a low metabolic activity, regardless of the presence or concentration of RGD. Whereas, on the intermediate stiffness substrate (56.83 KPa), crosslinked with 100 mM CaCl_2_, we can see a significant response to the presence of RGD on the cell metabolism, yet the response is not concentration dependent. However, on the stiffer substrate (114.33 KPa), crosslinked with 300 mM CaCl_2_, we can see a significant response to the presence of RGD on the cell metabolism which is concentration dependent. In addition, no significant difference on the cell metabolic activity was observed among substrates without RGD regardless of their stiffness (Fig. 8).

**Figure 8:**
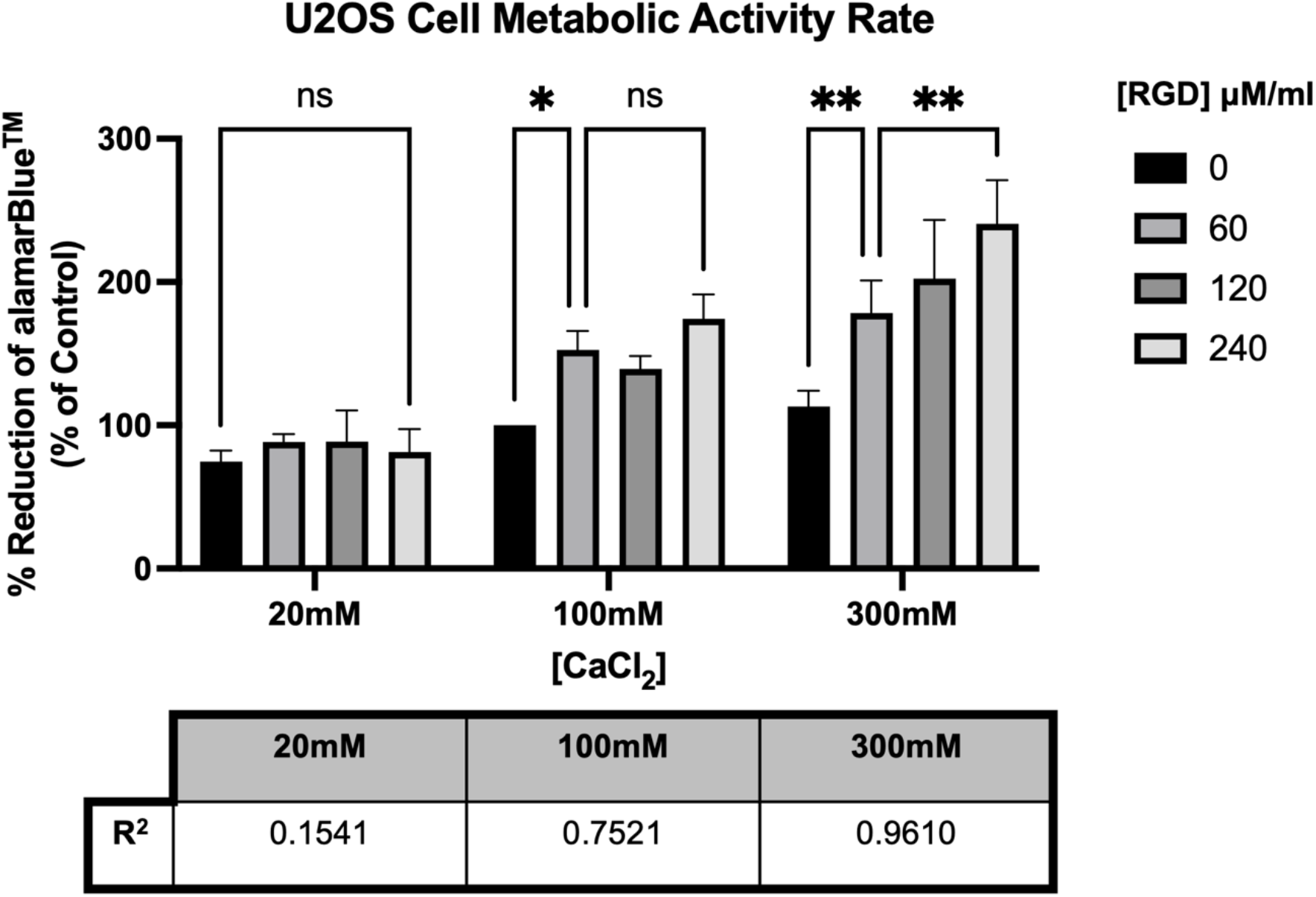
U2OS cells metabolic activity rate. Quantification of the % of alamarBlue^TM^ reduction of U2OS cells on 6% alginate + 2% gelatin, hydrogel H1, crosslinked with 20, 100 or 300 mM CaCl_2_ and coated with either 0, 60, 120 or 240 µM of RGD/mL on day 7 of cell culture. Calculations were done as percentage of the control (hydrogel crosslinked with 100 mM CaCl_2_ without RGD was considered 100%) (n= 4, Two-way ANOVA, p*<0.05, p**<0.01, ns: not significant). R-squared table of the correlation between the hydrogel stiffness and RGD content present on the alginate -gelatin hydrogel on U2OS metabolic response.

## 4. DISCUSSION

One of the most prominent issues related to the use of hydrogels on 3D (bio)printing is the poor mechanical properties of the scaffolds leading to their instability and complete degradation, thus limiting their use in basic research. To determine a suitable alginate-gelatin hydrogel for studying bone formation *in vitro*, four different formulations of hydrogel were tested for swelling and degradation rates during 28 days under cell culture conditions. All the hydrogels were prepared with 6% alginate to increase stiffness and yet allow a good printability in fluid-phase. H1 and H2 were prepared using MES buffer as a solvent whereas H3 and H4 were prepared with the complete cell culture medium RPMI 1640 supplemented with 10% FBS. Both solvents are widely used on hydrogel preparations for (bio)printing, but our results showed that MES buffer provided better stability for alginate-gelatin hydrogel in comparison to the RPMI medium. To increase cell attachment sites provided by gelatin, H2 and H4 were prepared with a higher gelatin concentration. However, increasing the gelatin content decreased the cell growth on the alginate-gelatin hydrogel. Gelatin behaves like fluid above 25°C (loss modulus G′′ > storage modulus G′) which leads to a decay on the mechanical strength of hydrogels containing more gelatin in their composition [41].

All the variables involved in the production of the (bio)printings are important for defining the mechanical, physical, chemical, and biological characteristics of the scaffolds. The type and concentration of the (bio)material(s), preparation protocol, crosslinking type, volume, and duration, for instance, are factors that can change the properties of the (bio)printings [42, 43]. In this study, we observed that the use of 6% alginate + 2% gelatin in 0.1 M MES buffer hydrogel crosslinked overnight with 5 ml of 100 mM CaCl_2_ allowed U2OS cell culture for at least 20 days presenting around 87% of cell viability on day 20 of cell culture. U2OS long-term cell culture showed an extensive presence of aggregates, multi-layered cell growth, and round-shaped monolayered cells. However, our results showed that it is possible to create alginate-gelatin substrates that drives spread or clustered cell growth by adding ECM or not or changing incubation time to coat the hydrogel with RGD, for instance. One of the mechanisms that determine the cell organization is epithelial-to-mesenchymal transition (EMT) which involves the downregulation of the expression of E-cadherins and upregulation of N-cadherins which decreases cell-to-cell and cell-to-matrix adherence and increases migration. This transition can be activated or deactivated by the cell response to the environment [44 - 47]. Having ways to drive cell behaviour, growth, metabolism, and organization is a valuable tool for the study of cancer biology, drug screenings or microtissue formation.

In this study, the NAIAD 3D bioprinter was used to produce fluid-phase printings within a cross-linkable aqueous support bath. Virtually all the chemical and biological reactions in the body need water to happen and obtaining (bio)printings in an aqueous phase allow us to develop protocols to enhance the scaffold’s biomimetics. In addition, our results showed that printing in fluid-phase allowed better printing fidelity and resolution at lower pressure and speed in comparison to air printing. Low pressure and speed decrease the shear stress, hence increases cell viability in bioprinting [48]. In this context, fluid-phase could possibly also be a way to improve bioinks 3D bioprinting. Moreover, according to Suntornnond *et al* (2016), higher speeds result in the increase of defects on the printed structures. Finally, Webb and Doyle (2017) suggested the use of the “Parameter Optimization Index” (POI) as a standard method to compare the quality of extrusion-based (bio)printings. The POI is calculated as: accuracy*modulus/pressure*printing width. Taking into consideration the comparison between fluid-phase and air printing in the context of the POI, the fluid-phase would show a higher POI score as it presents higher accuracy within lower pressures resulting in smaller printing strand width in comparison to air printing under the same conditions. Analyzing the evidence described in the literature together with our results, leads us to believe that the fluid-phase is not only a way to pattern the hydrogels during the printing process, but is also a way to optimize the printability parameters of the hydrogels (bio)printing within low pressures and speeds.

To date, there are few types of freeform or imbedding 3D (bio)printing described in the literature. Most of the techniques use the support bath to print aqueous phase solutions or soft hydrogels. Taking into consideration that a considerable part of human tissues is too soft to be accurately reproduced by extrusion-based air 3D bioprinting, we believe that the freeform printing is in the right direction to recreate human tissues *in vitro*. In this work we demonstrated the use of the support bath containing cross-link solution to tailor the hydrogel mechanical properties and biological pattern in a fluid-phase. Firstly, we showed that it is possible to add different ECM molecules to alginate-gelatin hydrogel using an aqueous cross-linkable solution bath via carbodiimides chemistry. Our fluid-phase method to incorporate ECM into the hydrogel showed to be efficient to make alginate-gelatin hydrogel attractive for cell colonization regardless of the seeding protocol. Moreover, we could compartmentalize and boost cell growth in pre-designed locations of the hydrogels. Other cell types that are also dependent on binding molecules for attaching and growing onto printed structures would also be suitable for compartmentalization by using the same methodology. Moreover, it could also be extrapolated to other biological reactions and addition of different molecules into printed structures such as others ECM, growth factors, signaling molecules, costimulatory molecules, etc.

Additionally, we showed that the cross-linkable support bath can also be used to tailor the mechanical properties of the alginate-gelatin hydrogel via fluid-phase methodology. By changing the cross-link concentration during the printing process, we produced substrates presenting different stiffness in the same printing. U2OS cells showed massive growth predominantly on the stiff part of the printing showing that we can also compartmentalize cell growth by patterning this aspect of the hydrogel. Tissues are not homogeneous mass with constant stiffness throughout their matrix. Therefore, this methodology could be valuable to print tissues with a wide range of stiffness, such as, for example, lungs, liver, and heart. Furthermore, it would also be advantageous to study the biology of diseases which involve changes in the stiffness of the healthy tissue. Our results also showed that U2OS cell number and metabolism rate were both increased on the stiff substrate (56.83 KPa), 3.26 and 2 folds, respectively, in comparison to the soft substrate (4.35 KPa). The difference in the cell number and metabolism ratios is probably due to the cell arrangement on the printings as U2OS form both monolayers and aggregates when growing on alginate-gelatin coated with RGD. Cells on spheroids formation present a lower metabolism in comparison to cells growing in monolayers [51]. Moreover, we observed an increase on cell death of cells growing on hydrogel crosslinked with 100 mM CaCl_2_ in comparison to cells growing on the same hydrogel crosslinked with 20 mM CaCl_2_ on day 1. High extracellular concentration of Ca^2+^ released from the alginate in the first hours of cell culture leads to an increase in cell death [52, 53]. However, this increase in cell death on the first day did not imply in a disruption of cell colonization of the printings, as on day 7 the hydrogels crosslinked with 100 mM CaCl_2_ showed greater cell growth in comparison to the hydrogels crosslinked with 20 mM CaCl_2_.

Finally, we showed how stiffness can affect the cell metabolic activity in response to the environmental concentration of RGD. In the present study, we observed that increasing the stiffness of the substrate can change the way the U2OS cells respond to the increasing concentration of RGD. The soft substrate (4.35 KPa) did not promote cell growth; therefore, cells did not respond to either the addition or the increasing of RGD. The intermediate stiffness (56.83 KPa) allowed massive cell growth over time and a significant increase on the cell metabolic activity with the presence of RGD, though did not show any difference on the cell metabolic activity in response to the increasing concentration of RGD. In these two circumstances, the response to RGD was most likely limited by the stiffness of the substrate as when the stiffness was increased to 114.33 KPA, we saw a concentration-dependent effect on the cell metabolic activity response to the extracellular concentration of RGD. This result illustrates the importance of tailoring multiparameter on 3D printings to provide cells with substrates more faithful to the *in vivo* systems to develop more accurate *in vitro* studies and more efficient drug screenings.

## 5. CONCLUSION

Tissues are dynamic environments with extensive diversification of biological molecules, chemical reactions, physical structures, and mechanical forces to support cell growth, differentiation, and activity. Reproducing the features of each one of the human tissues *in vitro* is a complex task, especially regarding the balance between the mechanical and the biological aspects of the (bio)printings. This study showed that 6% alginate + 2% gelatin hydrogel added with RGD is a potential substrate to grow bone *in vitro*. Additionally, the fluid-phase 3D printing showed to be a promising tool to design and pattern hydrogels. Overall, this methodology increased the shape fidelity and resolution of the printed structures within low pressures. Moreover, we could guide and compartmentalize U2OS cell growth to pre-designed locations in the printings by using the fluid-phase 3D printing methodology to tailor the hydrogel. Finally, U2OS cells showed a synergistic effect on the cell metabolic activity promoted by the customization of stiffness and ECM availability showcasing the importance of tailoring the substrate to guide cell growth and behaviour *in vitro*.

## ACKNOWLEDGEMENT

The authors acknowledge to Krutika Singh, PhD researcher affiliated to the School of Chemistry at University College Dublin, for her invaluable help with the rheological tests.

